# S-CAP extends clinical-grade pathogenicity prediction to genetic variants that affect RNA splicing

**DOI:** 10.1101/343749

**Authors:** Karthik A. Jagadeesh, Joseph M. Paggi, James S. Ye, Peter D. Stenson, David N. Cooper, Jonathan A. Bernstein, Gill Bejerano

## Abstract

There are over 15,000 known variants that cause human inherited disease by disrupting RNA splicing. While several *in silico* methods such as CADD, EIGEN and LINSIGHT are commonly used to predict the pathogenicity of noncoding variants, we introduce S-CAP, a tool developed specially for splicing which is better able to effectively distinguish pathogenic splicing-relevant variants from benign variants. S-CAP is a novel splicing pathogenicity predictor that reduces the number of splicing-relevant variants of uncertain significance in patient exomes by 41%, a nearly 3-fold improvement over existing noncoding pathogenicity measures while correctly classifying known pathogenic splicing-relevant variants with a clinical-grade 95% sensitivity.

## Introduction

Genomic sequencing, and in particular exome sequencing, is revolutionizing the diagnosis of Mendelian disease^1–4^, with over 5,000 genetic diseases already successfully mapped to over 3,000 genes^5^. Sifting through patient exomes in search of a causal variant is a time consuming process that often focuses on the coding sequence (CDS) of genes^6,7^. Powerful pathogenicity meta-predictors such as M-CAP^8^ integrate multiple primary predictors such as SIFT^9^, Polyphen-2^10^ and CADD^11^ with cross-species sequence conservation features to offer accurate clinical grade predictions capable of missing only a tiny fraction of known coding pathogenic mutations^12^. Achieving high sensitivity is important because reducing the size of the candidate list of variants of uncertain significance (VUS) is futile if the pathogenic variant itself is incorrectly classified as benign^13^. M-CAP is commonly used by clinicians to prioritize nonsynonymous variants^14,15^, so that they can effectively consider first the variants that are most likely to yield a diagnosis.

However, with estimates that 70% of patient cases remain undiagnosed^16,17^, there is value in looking beyond the CDS itself. Splicing is a complex and crucial step of gene expression, wherein vast sections of RNA, called introns, are removed from a pre-messenger RNA and the remaining RNA, called exons, are joined together to form the mature messenger RNA (mRNA). Changes in splicing induced by genetic variants can have severe impacts on the protein coding potential of an mRNA, such as the exclusion of an entire exon^18^ or a frameshift^19,20^ induced by creation of a new splice site^21^, among other effects^22–24^. Exome sequencing typically captures sequence information up to 50 base pairs past exon boundaries into each adjacent intron^25^. This region covers a broad class of splicing-relevant variants: those that disrupt existing splice sites or exonic and proximal intronic splicing regulators, such as the branch point, and some that create new splice sites.

Indeed, there are over 15,000 known Mendelian disease causing variants that impact the gene product through RNA splicing^26,27^. At the same time, a typical singleton patient’s exome contains over 500 splicing-relevant variants of uncertain significance (VUS)^28^. Existing tools such as CADD^11^, EIGEN^29^ and LINSIGHT^30^ tackle a broad spectrum of non-coding variants, but in doing so they do not effectively predict splicing-relevant variant pathogenicity or provide clinical grade assurances of minimizing the false prediction of known pathogenic splicing-relevant variants as benign. Generic methods ignore the rich literature characterizing mRNA splicing and predicting associated molecular phenotypes, such as the percentage-spliced-in of exons^31–34^. These findings and methods provide invaluable insight into the potential of a variant to disrupt splicing. However, the splicing literature does not tell the whole story either, as predicting molecular phenotypes is a fundamentally different task from predicting if a variant will cause a disease. For a variant to cause a disease, the variant must disrupt normal splicing in one or more relevant tissues *and* the induced change in mRNA phenotype must be pathogenic. Similar obstacles are met in the few cases where clinicians attempt to identify patient genetic variants that disrupt splicing by performing a costly and time consuming RNA-seq experiment in an accessible, but not necessarily disease relevant, cell population^35,36^.

We introduce S-CAP (**S**plicing **C**linically **A**pplicable **P**athogenicity), a machine learning tool that integrates knowledge of splicing with measures of variant, exon and gene importance into a splicing-specific pathogenicity score. We evaluate S-CAP at the high sensitivity required in clinical settings and show that it far outperforms existing non-coding pathogenicity scores, as well as tools focused solely on identifying synonymous variants that disrupt splicing. S-CAP will allow clinicians to consider a broad class of splicing-relevant variants, resulting in the diagnosis of more patients suffering from Mendelian disease.

## Results

We developed a machine-learning framework to model and evaluate splicing-relevant variant pathogenicity. We analyzed the positional distribution and potential functional effects of variants near splice sites and defined 6 genomic regions that display distinct mutation rates and functional effects. We trained 6 models (1 per region) to predict splicing variant pathogenicity. This involved building (i) a labeled dataset of known pathogenic and benign splicing-relevant variants, (ii) a set of features to help discriminate between the pathogenic and benign variants in each region and (iii) a learning algorithm to identify patterns in the features and to distinguish between variants in the two classes. Finally, we evaluated our models on a set of known pathogenic and benign variants, as well as on real patients with various diseases caused by splicing-relevant variants.

### The landscape of splice region variation

To build a dataset of labeled splicing-relevant variants, we considered variant pathogenicity, semantic effect and population frequency. We start by taking the union of 109,279 pathogenic single nucleotide variants (SNVs) from the Human Gene Mutation Database^26^ (HGMD) and 25,793 pathogenic SNVs from ClinVar^37^ to get a total of 114,382 unique pathogenic variants. We curated 15,833,389 benign SNVs observed in controls from the gnomAD database^38^ who generally do not suffer from a Mendelian disease. To identify a subset of variants with a likely splicing semantic effect, these sets were filtered to the ‘splicing region’, all synonymous or intronic variants within 50 base pairs^25,39^ of an exon boundary (we justify the choice of ‘splicing region’ below). Removing nonsynonymous and loss of function (stop gain or stop loss) variants ensured that our model was trained and evaluated nearly exclusively on splicing-relevant variants. To avoid any ambiguities, the benign SNVs were further filtered to remove any variant labelled pathogenic in HGMD or ClinVar. This resulted in 14,938 splicing-relevant pathogenic variants and 7,027,609 splicing-relevant benign variants. Then a frequency filter, based on the ACMG guidelines^13^ that suggest clinicians consider common (> 1% frequency) variants as definitively benign, was applied to both sets yielding 14,838 rare splicing-relevant pathogenic variants and 6,760,450 rare splicing-relevant benign variants. In support of this ACMG guideline, we note that only 100 of 14,938 (0.67%) known pathogenic variants in the splicing region are common in the general population (see Methods).

To substantiate that nearly all pathogenic ‘splicing region’ variants do, in fact, disrupt splicing, we developed a simple model to assign a putative effect of variants on splicing. Variants can affect splicing by (1) create a cryptic splice site (2) disrupt an existing splice site or (3) disrupt an existing branchpoint, a mechanistically essential sequence motif generally located 18 to 45 base pairs upstream of each 3’SS^40^. We used the existing tools MaxEntScan^41^ and LaBranchoR^42^, which predict the strength of splice site sequences and branchpoint sequences, respectively. We denoted a variant as creating a cryptic splice site if the variant creates a splice site with a MaxEntScan score at least as high as the score for the reference splice site (with the variant included), disrupting a splice site if it greatly reduces the MaxEntScan score of the reference splice site, and disrupting a branchpoint if it has a low LaBranchoR *in silico* mutagenesis score. In cases where a variant had multiple putative effects, e.g. creating a cryptic splice site and disrupting an existing splice site, we resolved to the most extreme effect (cryptic splice site creation > splice site disruption > branchpoint disruption). We found that 97% of pathogenic splicing-region variants are predicted to have an effect on splicing, as opposed to only 18% of likely benign variants (**Fig. 1a-b**). We found that pathogenic variants are enriched at positions where the mechanistically essential U2 snRNA, U2AF, and U1 snRNA bind^43^ (**Fig. 1b**). Conversely, variants occurring in the general population are biased away from these high information content positions^44^ (**Fig. 1a**).

**Figure 1.**
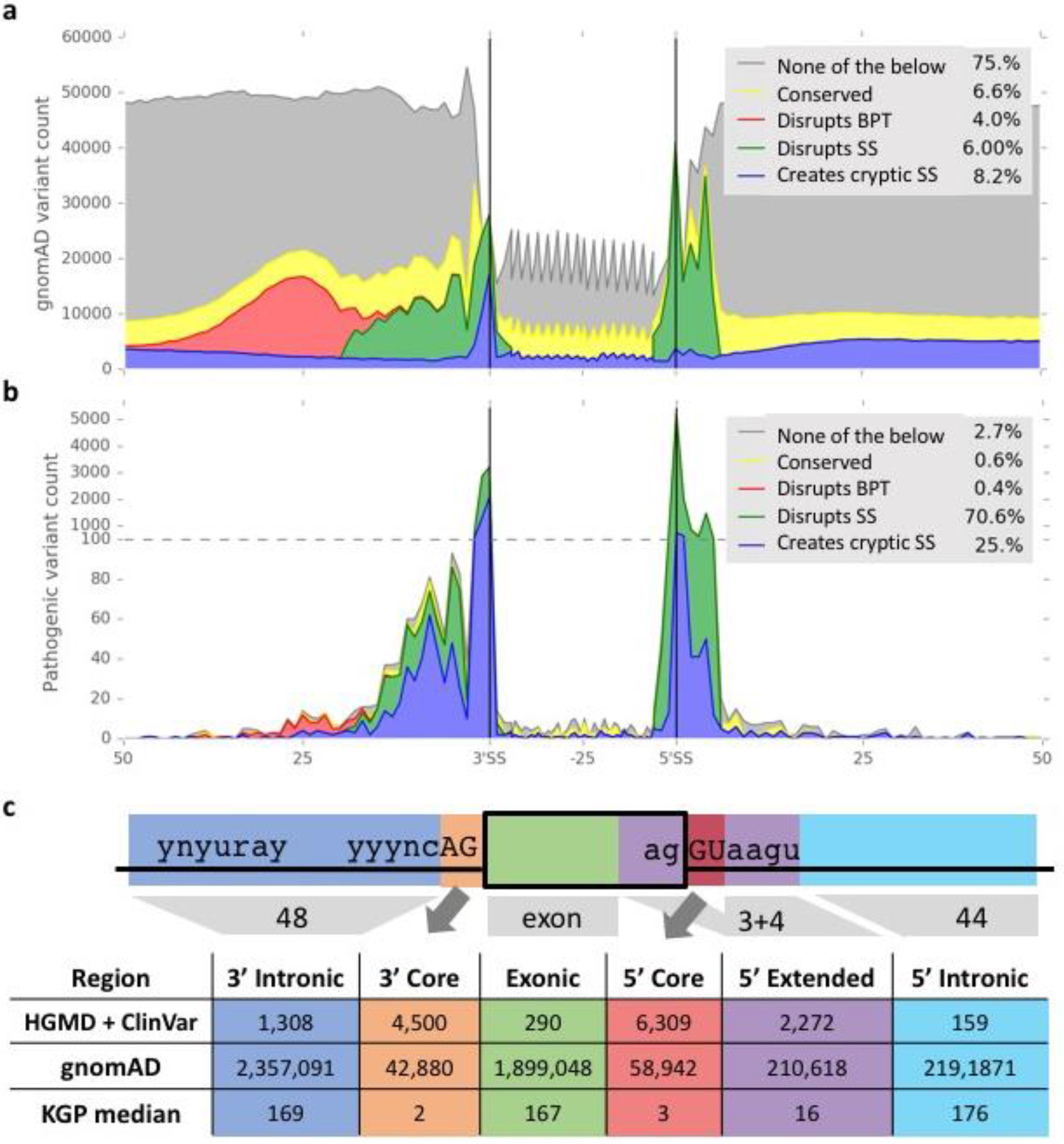
Distribution of rare, noncoding variants in the splicing region. We built two sets of rare variants near exon boundaries that have no effect on the annotated coding sequence: (A) a set of likely benign variants from the Genome Aggregation Database (gnomAD) and (B) a set of putatively pathogenic variants from the Human Gene Mutation Database (HGMD) and ClinVar. We developed a simple model to ascertain the effect of these variants on splicing, using MaxEntScan to assess if a variant created a cryptic splice site (SS) or disrupted the reference splice site, and LaBranchoR *in silico* mutagenesis scores to assess if a variant disrupted a branchpoint (BPT). In (A) and (B), we plot the aggregate counts of each variant set as a function of position relative to the nearest splice site, colored by their putative effect. Nearly 97% of pathogenic variants in the splicing region (as defined in the text) are predicted to have a putative deleterious effect on splicing, as compared to only 18% of benign variants. (C) We split the variants into different regions with largely homogenous function. The majority of known pathogenic variants (and the easiest ones to detect) are found in the 3’ and 5’ core splice site regions, whereas the majority of benign variants are found in the 3’ intronic, exonic and 5’ intronic regions. In a typical (median) individual from the 1000 genomes project (KGP), the distribution of variants is similar to the distribution of gnomAD benign variants.

### Region specific models to increase performance and alleviate ascertainment bias

In order to effectively capture these position specific patterns, we separated the variants into 6 regions relative to the splice sites with generally homogenous function and built a separate model for each region (**Supplementary Fig. 1**). Specifically, we grouped variants occurring in the obligate 5’ GT (5’ core) and 3’ AG (3’ core) dinucleotides, intronic variants upstream of a 3’ss (3’ intronic), variants lying in the canonical U1 snRNA binding site, excluding the core 5’ss (5’ extended), intronic variants downstream of a 5’ss (5’ intronic), and synonymous variants within the protein coding gene (exonic) (**Fig. 1c**).

Core splice site variants are well known as having a large functional effect and are readily identified, a fact that has likely led to an overrepresentation of core splice site variants in pathogenic variant databases. Around 73% of known splicing region pathogenic variants occur within the core splice sites (**Fig. 1c**). Although this is consistent with the mechanistic importance of these positions, in unbiased studies of splicing quantitative trait loci, it is generally found that fewer than 1% of splicing-relevant variants are found to be located in core splice sites^45,46^ and in a recent study employing RNA-seq data to identify splicing-relevant variants resulting in Mendelian disease only 2 of 6 (33%) causal variants in the splicing region were in core splice sites^35^. If left unaddressed, this bias would allow a classifier to have strong test set performance (by calling most core splice site variants pathogenic and others benign), but would often miss non-core splice site pathogenic variants. Separating variants by position allows us to *guarantee* that pathogenic variants are rarely misclassified as benign regardless of genomic region, thereby ensuring an overall low false negative rate irrespective of ascertainment biases present in annotated data.

### S-CAP features

We curated existing metrics and developed several novel features to help distinguish between pathogenic and benign variants within the splicing region (**Supplementary Table 1**). The set consists of chromosome, gene, exon, and variant level features. At the **chromosome level**, 3 binary features distinguish between variants found on chromosome X, chromosome Y and the autosomes. Variants on the X chromosome present an important subset since in males, a hemizygous X chromosome variant inducing loss of function results in no viable gene product. Consistent with this intuition, pathogenic variants are highly enriched on the X chromosome as compared to the autosomes (7.11 fold enrichment, p < 10^−140^ by two-sided Fisher’s Exact Test). At the **gene level**, pLI^44^, RVIS^47^, and a haploinsufficiency score^48^ help to measure the likelihood that a given gene is a ‘disease gene’. At the **exon level**, exon length, exon length modulo 3, reference splice site strengths, an existing regional constraint score^49^ and the exon sequence similarity between hg19 and 99 species from the UCSC 100way alignment serve to assist in distinguishing critical exons from those that may be safely excluded. Additionally, we developed a novel splice site constraint score to measure the fragility and tolerance of each exon to splice site mutations (see Online Methods). At the **variant level**, CADD^11^ measures pathogenicity based on functional data annotations, LINSIGHT^30^ measures variants’ fitness effect through functional data and molecular evolution and SPIDEX^33^ was incorporated to measure the impact of a variant on exon inclusion. PhyloP^50^ and PhastCons^51^ scores from the multiz46way and multiz100way alignments^52^ measure the evolutionary importance of the affected base across primate species, placental mammals and all vertebrates. We also included a feature to capture the change in 3-mer content induced by a variant^53^. Additionally, we included region-specific features, such as a branchpoint disruption term for the 3’ intronic region^42^ and a 5’ cryptic splice site creation term for the 5’ intronic region (see Online Methods for a complete description of features).

### The machine learning algorithm

Similar to the M-CAP classifier^8^ for nonsynonymous SNVs, S-CAP is built using a gradient boosting tree classifier, a highly effective machine learning model^54^. This model iteratively builds decision trees, where each tree is picked to correct the most cases that were misclassified in the previous step. The final classifier is a linear combination of each of the previously derived decision trees (see Online Methods for details).

### S-CAP consistently outperforms existing pathogenicity scores

Each of the 6 regions (described above) contains a set of pathogenic and benign variants (**Fig. 1c**), which were used to train 6 separate models. We performed 5-fold cross-validation and selected the median performing model as the final model for each region. S-CAP was evaluated against the most popular existing methods that score splicing-relevant variants: CADD, SPIDEX, LINSIGHT and EIGEN. The 6 S-CAP models outperformed all existing methods in all regions resulting in up to a 26.6% improvement in the AUC over the next best performing model (**Fig. 2**). S-CAP performance ranged from achieving an AUC of 0.804 in the 3’ core region (**Fig. 2b**) to achieving an AUC of 0.953 in the 3’ intronic region (**Fig. 2a**). No existing method consistently outperforms the other existing metrics across all splicing regions (**Fig. 2**).

**Figure 2.**
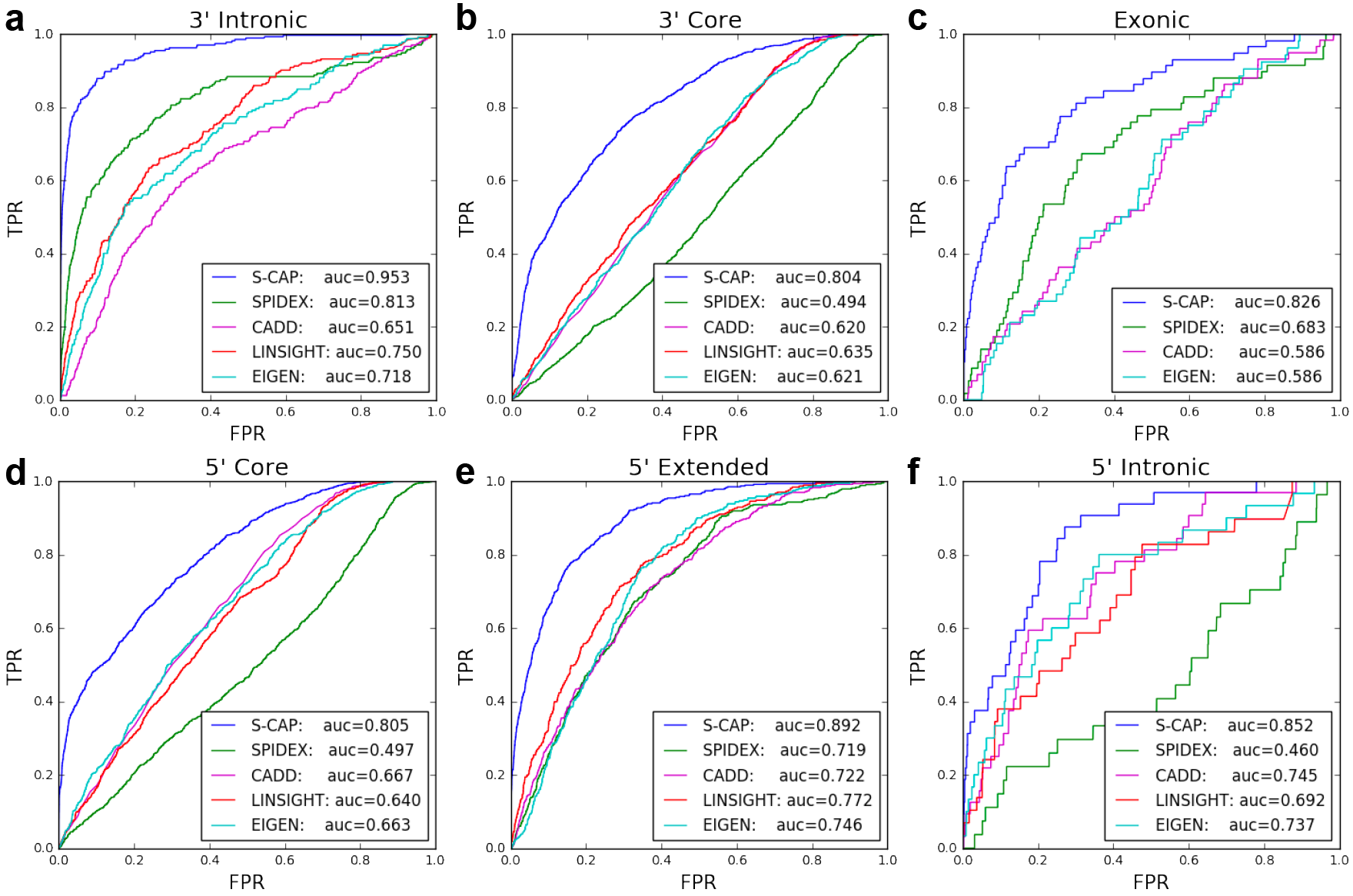
Overall performance per region for splicing pathogenicity classification. For each method, 1000 threshold points were determined by evenly spanning the range from the minimum to the maximum score observed for the method. A true positive rate and false positive rate were determined for each threshold value and used to build the receiver operating characteristic (ROC) curve. We compared to the next best method in each region and found that S-CAP performs 17.2% better in the 3’ intronic region (A), 26.6% better in the 3’ core sites (B), 20.9% better in the exonic region (C), 20.7% better in the 5’ core sites (D), 15.5% better in the 5’ extended region (E) and 14.4% better in the 5’ intronic region (F). S-CAP outperforms existing metrics in all regions whereas none of the existing methods consistently outperforms the others.

To ensure a fair comparison, S-CAP performance on exonic variants was also independently compared against MutPred Splice^53^, a tool focused on scoring only splicing-relevant synonymous variants, and found superior in both AUC and especially hsr-AUC (see Online Methods and **Supplementary Fig. 2**).

### Clinically relevant threshold maintains high sensitivity

As previously shown in M-CAP^8^, it is important to tune thresholds for clinical settings so that fewer than 5% of known pathogenic variants are misclassified. Neither LINSIGHT, EIGEN or SPIDEX provides a default threshold to consider a variant as pathogenic. As a result, it is difficult to use these methods for variant pathogenicity classification. CADD provides a threshold but, at the author-recommended default threshold, virtually all pathogenic splicing mutations outside the two core splice sites are discarded and incorrectly classified as benign (**Table 1**). Similarly, MutPred Splice provides a default threshold, but over 48% of known pathogenic exonic variants are misclassified at this threshold. We generated a high sensitivity threshold for all metrics in each region by finding the lowest threshold that results in the correct classification of at least 95% of a test set of pathogenic variants from that region (**Table 2**).

**Table 1.** Misclassification Rate (Low Is Good) Of Existing Metrics At Author-Recommended Thresholds. The two tools that provide classification thresholds, misclassify 48%-99% of known pathogenic variants in different regions. Other tools do not provide a threshold.

|  | Method |  |  |
| --- | --- | --- | --- |
| Region | CADD | LINSIGHT,EIGEN,SPIDEX | Mutpred Splice |
|  | Author's Thresholds |  |  |
|  | ≥ 20 | N/A | ≥ 0.6 |
| 3' intronic | 99% | N/A | N/A |
| 3' core | 4% | N/A | N/A |
| exonic | 98% | N/A | 48% |
| 5'core | 2% | N/A | N/A |
| 5' extended | 96% | N/A | N/A |
| 5' intronic | 97% | N/A | N/A |

### S-CAP is the best performer at clinically relevant thresholds

Moving into the high sensitivity domain, at or above a 95% true positive rate, S-CAP’s performance (**Supplementary Fig. 3a-f**) ranged from achieving an hsr-AUC of 0.186 in the exonic region (**Supplementary Fig. 3c**) to an hsr-AUC of 0.549 in the 3’ intronic region (**Supplementary Fig. 3a**). Similarly, S-CAP outperforms MutPred Splice on an independent test set of exonic variants (**Supplementary Fig. 2b**). Overall, S-CAP improves on existing metrics in all 6 splicing regions when focused on the high sensitivity domain, and no existing method consistently outperforms the others.

### Different patterns observed for recessive and dominant variants

Recessive and dominant diseases are associated with different selective pressure on alleles, which we hypothesized would result in different feature importance and thresholds for determining variant pathogenicity. To address this complexity, we developed separate classifiers for dominant (heterozygous in patient) and recessive (homozygous or compound heterozygous in patient) at the gene level. In patients, we are given whether each variant appears in the heterozygous or homozygous state. However, our training data did not provide dominant and recessive labels, so we developed a framework for predicting this information from a control population for the sake of model training (see Methods).

Tagging 3’ and 5’ core variants as dominant or recessive resulted in improved performance **Supplementary Fig. 4a-h**) compared to models without these tags (**Fig. 2b,d**). For 3’ core variants, we built a model that improved upon an AUC of 0.804 and hsr-AUC of 0.257 from the original 3’ Core model to an AUC of 0.805 and 0.890 (**Supplementary Fig. 4c,d**) and hsr-AUC of 0.296 and 0.454 (**Supplementary Fig. 4g,h**) when tested on just dominant or recessive variants, respectively. Similarly, for 5’ core variants, we built a model that improves upon an AUC of 0.805 and hsr-AUC of 0.291 from the original 5’ core model to an AUC of 0.779 and 0.880 (**Supplementary Fig. 4a,b**) and hsr-AUC of 0.222 and 0.518 (**Supplementary Fig. 4e,f**) when testing on just dominant or recessive variants, respectively. The S-CAP model to be used on patients takes advantage of these split dominant and recessive models in the core splice site regions.

### S-CAP eliminates the most VUS in patient exomes

Resources like S-CAP are developed on large sets of benign and pathogenic variants but ultimately are used to help with the interpretation of VUS in individual patients (**Fig. 1c**). To demonstrate the practical utility of S-CAP, we evaluated S-CAP and each of the comparison methods on 14 patients with Mendelian diseases caused by splice-altering mutations. After applying the standard allele frequency filter of ≤ 1%, a typical individual has on average a total of 533 rare variants within the splicing region (**Fig. 1c**). Typically, ^~^32% of variants were observed in each of the 5’ intronic, exonic and 3’ intronic regions. 3% were in the 5’ extended regions and 0-3 variants were in each of the 3’ core and 5’ core regions (**Fig. 1c**). Existing methods, CADD, LINSIGHT, EIGEN and SPIDEX, using the 95% sensitivity thresholds (**Table 2**), perform comparably when applied to all VUS in the splicing region of an individual patient and on average reduce the number of VUS by 4% - 15%. By contrast, S-CAP is more powerful, reducing the number of VUS in the splicing region for an individual patient by 31-46% (**Fig. 3c**), while confidently retaining the pathogenic variant for further detailed analysis (**Table 3**). For a larger sample size, we also evaluated each method on all (n=2054) individuals in the 1000 Genomes Project^28^, which can conceptually be thought of as Mendelian disease patients with their pathogenic variants removed (see **Fig. 1c**). The observed fraction of VUS reduced on average per 1000 Genomes Project individual is consistent with the performance observed when applied to patients (**Fig. 3c-d**). Specifically, S-CAP is nearly three times as powerful as existing methods and on average results in a 41% reduction of the VUS within the splicing region of a given individual.

**Table 2.** High sensitivity thresholds after recalibrating each method. We retuned each existing method for each of Figure 1’s six regions by finding the smallest threshold that resulted in the correct classification of 95% of pathogenic variants from that region. S-CAP thresholds for all regions (including the dominant and recessive modes) are included in the last column. By definition with the high sensitivity thresholds each method will misclassify at most 5% of known pathogenic mutations, but their effectiveness at correctly classifying benign variants and reducing the number of patient VUS will vary greatly.

| region | CADD | LINSIGHT | EIGEN | SPIDEX | MutPred<br>Splice | S-CAP |
| --- | --- | --- | --- | --- | --- | --- |
| 3' intronic | ≥ 0.709 | ≥ 0.048 | ≥ 3.57 | ≤ -1.45 | N/A | ≥ 0.005 |
| 3' core | ≥ 21.50 | ≥ 0.767 | ≥ 7.25 | ≤ 1.14 | N/A | Dom.: ≥ 0.031 |
|  |  |  |  |  |  | Rec.: ≥ 0.144 |
| Exonic | ≥ 0.061 | ≥ 0.086 | ≥ 3.78 | ≤ -2.31 | ≥ 0.090 | ≥ 0.012 |
| 5' core | ≥ 22.6 | ≥ 0.795 | ≥ 8.381 | ≤ 1.65 | N/A | Dom.: ≥ 0.032 |
|  |  |  |  |  |  | Rec.: ≥ 0.357 |
| 5' extended | ≥ 7.423 | ≥ 0.211 | ≥ 13.80 | ≤ -0.910 | N/A | ≥ 0.003 |
| 5' intronic | ≥ 0.852 | ≥ 0.047 | ≥ 0.847 | ≤ -1.636 | N/A | ≥ 0.004 |

**Table 3.**
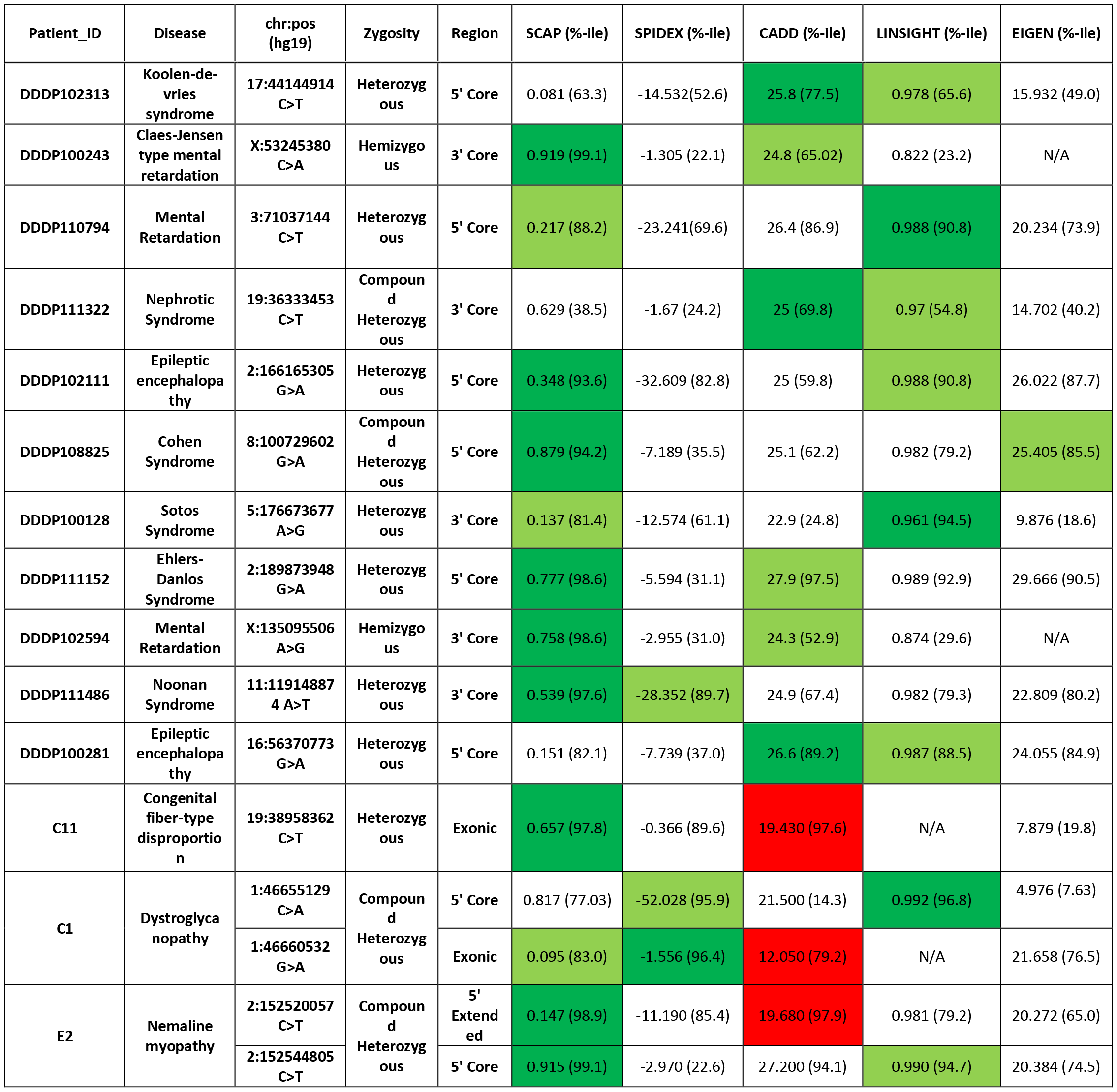
Causative variants in each patient and pathogenicity predictions. Each row describes a single patient, their underlying disease, causative variant and its zygosity, the region of the genome in which it is located, and the score and percentile each method assigns to the variant. The percentile was computed by measuring the fraction of variants in the same region with a score less than the score assigned to the causative variant. The 99^th^ percentile denotes that 99% of variants in that region have a score less-pathogenic than the one we are observing thereby indicating that the variant is considered to be highly pathogenic. Dark and light green filled entries have the highest and second highest percentile scores for the given variant, respectively. Red entries highlight patients where the causative variant would have been classified as benign using the author-recommended thresholds.

**Figure 3.**
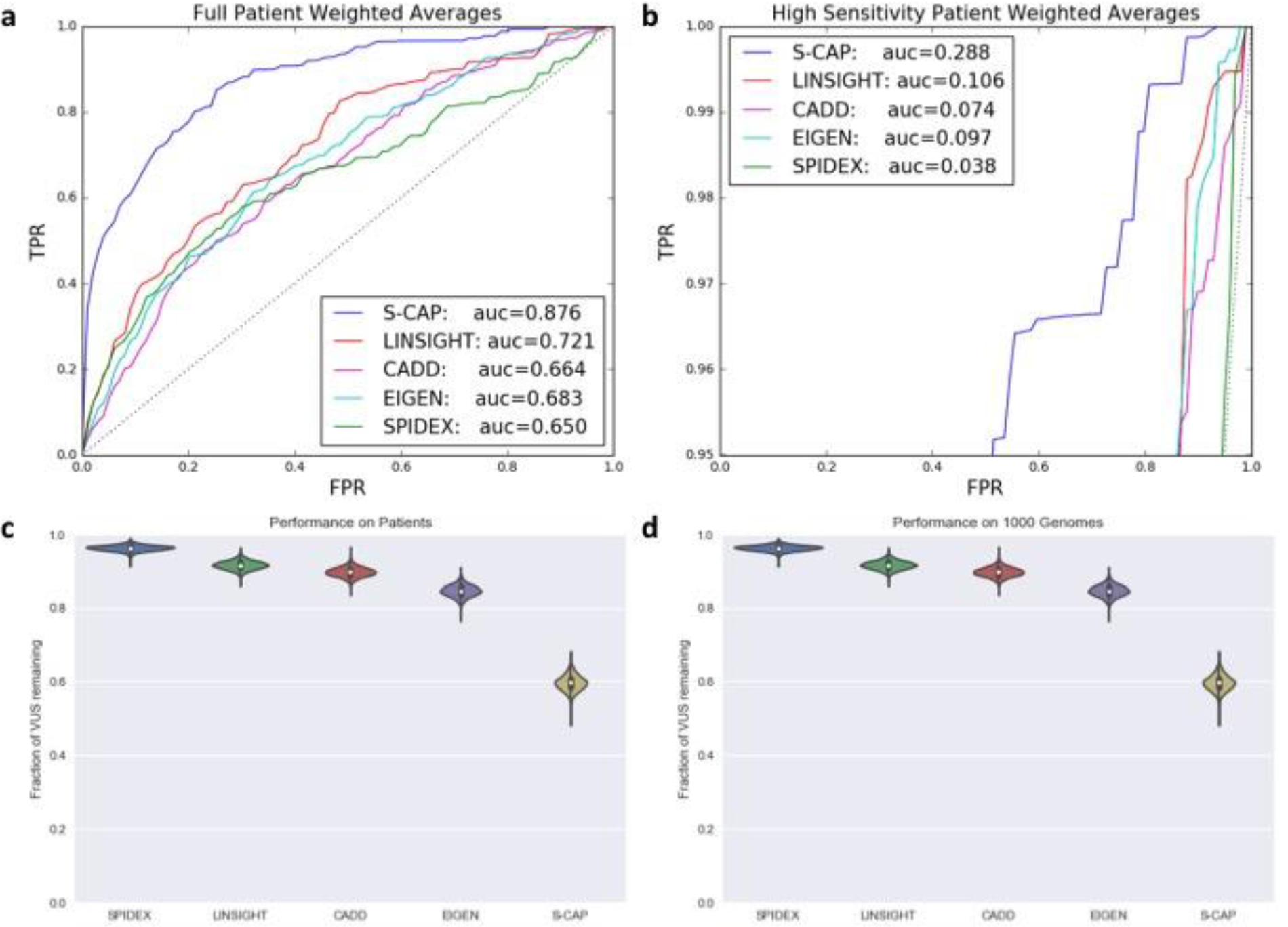
Overall performance on patient data. We take a weighted sum of the AUC from each of the 6 regions based on the distribution of variants seen in a typical individual to form an overall receiver operating characteristic (ROC) curve representative of the overall performance expected on patients. (A) S-CAP improves on the next best method by 21.5% in overall AUC and (B) and by 172% in the hsr-AUC. (C) S-CAP reduces the number of splicing related variants of uncertain significance (VUS) from patient exomes by 40% while maintaining the pathogenic variants with 95% sensitivity. At the same sensitivity requirement, existing methods reduce the VUS by only 4% - 15%. (D) We observe a similar reduction in VUS over all (n=2054) Thousand Genomes Project individuals, which conceptually only differ from Mendelian disease patients by up to 2 mutations.

## Discussion

Variants affecting splicing comprise the second largest category of known pathogenic mutations^27^. A broad class of potential splicing-relevant variants are already being captured by exome sequencing, yet clinicians are less able to interpret these variants for lack of proper interpretation tools. Here we address this problem by developing S-CAP, the first clinically applicable pathogenicity predictor dedicated exclusively to splicing-relevant variants.

In order for an *in silico* pathogenicity predictor to be useful in the clinic, it needs to be easy to use, carefully evaluated at high sensitivity and confidently remove a substantial fraction of benign variants. Existing noncoding variant tools are not easy to use for splicing variants because their performance is strongly dependent upon position relative to splice sites and no single method consistently outperforms the others. This means that employing existing methods, a clinician would have to consult multiple pathogenicity scores for variants in different regions. Furthermore, none of the existing methods have been carefully evaluated at clinical-grade sensitivity and most methods give no guidance about what cutoff should be used or result in the misclassification of an unacceptably high number of pathogenic variants (**Table 1**). After carefully evaluating the existing methods, we found that none of them confidently removed a considerable fraction of benign variants (**Fig. 3**). For example, after we retuned it (**Table 2**), SPIDEX performed well on variants in the intronic bins, but was close to random at predicting the pathogenicity of core splice site variants. CADD, only after we retune it, (**Table 1**,**2**) performed well on core and exonic variants but poorly on intronic variants (**Fig. 2**). S-CAP, addresses these important issues, as it consistently outperforms existing methods across all the regions, its performance has been carefully evaluated at clinical-grade sensitivity, and it removes close to three times as many benign variants as any other method (**Fig. 3**).

Central to the design of S-CAP is the use of region-specific models to alleviate the effects of ascertainment biases in curated pathogenic variant databases. Curated pathogenic variant databases contain invaluable information about the properties of pathogenic variants, but they also over-represent variants in known disease genes and in association with easily identifiable features. Of particular concern in splicing pathogenicity prediction is the inflated number of variants in core splice sites, which exists because they are easily recognized and have well established molecular consequences. If left unaddressed, this bias in the labeled pathogenic data would lead to unrealistic model performance, as a model could achieve relatively high test set performance simply by predicting that all core splice site variants are pathogenic and all others are benign. However, in the clinic, such a model would incorrectly classify pathogenic, non-core splice site variants as benign at an unacceptably high rate. Introducing separate models for each region alleviates this concern since each model’s performance is evaluated using data from the same region, thereby assuring high sensitivity irrespective of the underlying positional distribution of pathogenic variants. Additionally, we accounted for the over-representation of pathogenic variants in known disease-associated genes by ensuring that variants from the same gene were never split between the training and evaluation sets (see Methods). This guaranteed that no gene level information was shared by features across folds.

The use of patient RNA-sequencing (RNA-seq) data to identify pathogenic variants that disrupt splicing is a growing and promising field^35,36,55^ to which we believe S-CAP is complementary. In fact, our model already includes scores from SPIDEX^33^, a deep learning model that was trained on tissue-specific RNA-seq data to predict the change in percent spliced-in (*ΔΨ*) of an exon given a variant. It would only be a small step to supplement these predicted *ΔΨ*s with experimentally measured *ΔΨ*s from RNA-seq experiments. Observing through *ΔΨ* RNA-seq bypasses the difficult problem of explicitly or implicitly predicting the effect of a given variant on splicing in a particular cell type, but this is only part of the problem. Whether predicted or measured, gene expression *Ψ* and values vary between cell contexts and time points and in many cases the most relevant cell population to sequence would not be clear^36,56^. Perhaps even more importantly, molecular phenotypes are multiple steps away from real phenotypic change, making it difficult to predict organismal pathogenicity from molecular phenotype^57^. These factors limit the direct applicability of observed or predicted *ΔΨ*s to cases where there is a good understanding of the relationship between a disease, the cell population it affects, and the set of potentially causative genes. Many of the features used by S-CAP, such as evolutionary conservation, implicitly integrate an allele’s importance over all cell populations and time points, complementing the strengths of RNA-seq based methods. Another direction for the integration of S-CAP and experimental methods is the use of cheap and fast site-directed sequencing in a diverse array of cell types to validate putatively pathogenic sites identified by S-CAP^58^.

Eventually, comprehensive analysis of the splicing region will become commonplace in clinical settings. This is currently a difficult task given the complexity of splicing and the difficulty in predicting whether a change in splicing will result in disease. There are over five hundred rare variants of uncertain significance per individual in the splicing region with no clear semantic effect outside of the core splice sites and most are not observed in control populations. S-CAP represents an important step towards effectively interpreting splicing variation, but we will have to continue to improve on these methods, learning more from RNA-seq experiments, to render this problem more tractable.

## Online Methods

### Variant Processing

#### Dataset of pathogenic and benign variants

Pathogenic variants were obtained from two manually curated databases: the Human Gene Mutation Database (HGMD) Professional version 2017.1 and ClinVar release 20170406. Only HGMD variants tagged as Disease Mutation (DM) and ClinVar variants with Pathogenic Clinical Significance were included in the final set of pathogenic variants. Benign variants were obtained by identifying variants observed in individuals from gnomAD^38^ r2.0.2.

#### Variant Annotation

ANNOVAR^59^ v527 was used to annotate variants with predicted effect on protein-coding genes using gene isoforms from Ensembl^60^ gene set version 75 for the hg19/GRCh37 assembly of the human genome. All coding isoforms were used where the transcript start and end sites were marked as complete and the coding span was a multiple of three.

#### Variant Filtering

All variants were filtered so as to only include rare variants in the splicing region that do not directly affect the protein coding sequence. Rare variants are defined to be variants with an allele frequency of 1% or less in all control populations and subpopulations in KGP phase 3, ExAC v0.3.1 and gnomAD r2.0.2. Variants in the splicing region that do not affect protein coding sequence are those determined to be synonymous or in the core splicing, extended splicing or intronic regions.

### Positional Subsetting of Variants

Variants at different positions relative to splice sites have different properties, such as distributions of sequence conservation and ratio of pathogenic variants to benign variants. To better understand the relationship between position relative to splice site and the mechanistic effects on splicing, we developed a simple model to assign a putative effect of variants on splicing. To identify variants that create cryptic splice sites, we scanned a window the width of the MaxEntScan^41^ input for the highest scoring splice site that is not the reference and reported the variant as creating a cryptic splice site if this highest scoring site was stronger than the reference (with the alternative allele included) and is made stronger by the variant. Variants that result in a decrease the MaxEntScan score of the reference splice site by 1 or more were determined to disrupt an existing splice sites. Finally, variants that have a LaBranchoR^42^ *in silico* mutagenesis score of less than −0.1 were determined to disrupt a branchpoint.

This led us to allocate variants into subsets based upon their position relative to splice sites and train separate models for each. Subsets were selected so as to group together functionally related positions. In total, we constructed 6 subsets (**Fig. 1**). We created subsets of 3’ and 5’ core (1 or 2 bases from the splice site), extended (1-2 bases into the exon on the 5’ side and 3 to 6 bases from the 5’ splice site), 3’ intronic (2 to 50 bases from the splice site), 5’ intronic (7-50 bases from the 5’ splice site) and exonic (inside the exon but outside the extended region) regions.

We associated each variant with a neighboring exon. For variants close to multiple exons, we attempted to assign the variant to the exon considered as having the highest chance of a pathogenic effect, which we inferred from the density of pathogenic variants in each bin. Specifically, we favored associations in the following order: core, 5’ extended, intronic, and exonic.

### Features

Our models utilized a diverse library of previously described and novel features (fig. S1). These can be divided into gene level, exon level, and variant level features as well as a few features that are specific to individual regions. Below, we introduce our novel features and describe how we curated previously described features.

#### Gene Level

We obtained **RVIS**^47^ scores from, the appropriately named, genic-intolerance.org (see URLs). We used the data in column “RVIS[pop_maf_0.05%(any)]” as a feature in our models. We obtained **pLI**^44^ scores from supplementary table 13 of the original publication on 7 April 2017. We used as a feature the column denoted as pLI. We obtained a **haploinsufficiency score**^48^ which is a probability of a gene being haploinsufficient directly from the publication page on the PLOS website in May 2017. We downloaded MPC, a recently proposed regional constraint score^49^ (see URLs).

#### Exon Level

For each exon, we created **splice site features** which measure the number of rare and common variants observed in gnomAD in the 5’ and 3’ core (0-2) and extended (2-6) regions. This was motivated by the desire to share information between functionally similar positions. We take this feature to represent the constraint on the exon’s splicing region in the human population. To avoid data leakage, when constructing this feature for a particular variant, we masked the effect of the variant itself on this score. Specifically, if a given 5’ss had one associated core variant, it was assigned a count of 0. If it had 2, both were assigned a count of 1. Additionally, we measured **exon identity** across vertebrates and found that the exon identities in many organisms were highly correlated. Principal components analysis (PCA) of the identity scores for all exons showed that 5 components explain the vast majority of variation in the data. To prevent overfitting, we included the original exon identities projected onto these first 5 principal components as features. We also included exon length and exon length mod 3 as features associated with each variant.

#### Variant Level

In order to consider the local sequence context of variants, we included **spectrum kernel features** representing the change in trinucleotide content induced by the variant^53^. For all 64 possible trinucleotides, we created a vector counting the number of occurrences in the alternative sequence and subtracted an equivalent vector for the reference sequence. The **MaxEntScan reference score, alternative score and difference** between the two were all used as features to quantify the strength of each exons’ reference and alternative 5’ and 3’ splice site. **SPIDEX** scores^33^ were downloaded directly from the Deep Genomics website on 7 April 2017. These scores can now be downloaded from Annovar (see URLs). Any variant not assigned a SPIDEX score was assigned a value of 0. We included 8 scores measuring evolutionary conservation, specifically, **PhyloP 46way vertebrates, placental mammals, primates, PhyloP 100way vertebrates, PhastCons 46way vertebrates, placental mammals, primates and the PhastCons 100way vertebrates**. Any base not annotated with a conservation score was assigned a value of 0. We also included the signed **distance to the 5’ss and 3’ss splice site of the associated exon** as features to measure base-pair importance.

#### Region Specific features

The 3’ intronic S-CAP model includes a **branchpoint feature** modeled using LaBranchoR, a bi-directional LSTM (long short-term memory) model trained on the genome sequence surrounding experimentally validated branchpoint sites^42^. Specifically, we used the *in silico* mutagenesis scores available online (see URLs) as a feature.

We explicitly represented the strength of **cryptic sites** created by each variant using MaxEntScan^41^. In 3’ intronic, 3’ core, and exonic bins, we included a 3’ cryptic splice site creation term and, in the exonic, 5’ extended, 5’ core, and 5’ intronic bins, we included a 5’ cryptic splice site creation term. For each variant, we scanned for the highest scoring splice site motif that overlaps the variant, excluding reference splice sites. We used as features the strength of the cryptic site, the change in the strength of the cryptic site induced by the variant, the distance from the reference splice site to the cryptic site and the difference in strength between the cryptic site and the reference splice site.

### Model Training and Testing

We performed 5-fold cross-validation to train and identify a generalizable S-CAP model. 5-fold cross-validation refers to splitting the data into 5 roughly equally sized parts (folds). All variants found in a single gene were included in the same fold to ensure that there was no leakage of feature information across the training and test sets. We then merged 4 of 5 sets to form a training dataset, trained the model on this training dataset and evaluated on the remaining fold to obtain an expected accuracy. We performed this process 5 times (each combination of 4 folds was merged together to form the training dataset) testing on the fold that was not included in the training dataset.

To train the S-CAP model, we used a Gradient Boosting Tree model implemented in the python 2.7.13 sklearn version 0.18.1 library and used the default parameters to reduce the chance of overfitting the model. After training 5 models during the cross-validation phase, we picked the median performing model as the final classifier. The ROC curves were built based on performance on the test set for this specific median model.

### Comparison Metrics

We sought to compare our performance to those of other methods used to infer the importance / pathogenicity of noncoding variants. We evaluated the performance of each of the methods below by using the output score directly as a pathogenicity score. For SPIDEX, we negated the score, as is consistent with large negative scores having a larger impact on function and constraint, respectively. We report performances for all subsets where the method reported scores for at least 50% of variants. Variants where a method did not report a score are excluded from evaluation.

**CADD**^11^ v1.3 scores were downloaded from the CADD website (see URLs). **SPIDEX**^33^ scores were downloaded directly from the Deep Genomics website on 7 April 2017. Any variant not assigned a SPIDEX score was assigned a default value of 0. **LINSIGHT**^30^ scores were downloaded directly from the LINISGHT website (see URLs) on 7 April 2017. Any variants not assigned a LINSIGHT score was defaulted to be 0. **Eigen v1.0 coding** and **Eigen v1.0 noncoding**^29^ Phred scores were downloaded from the Eigen website (see URLs). **MutPredSplice**^53^ score were downloaded from the MutPredSplice website (see URLs).

As MutPred Splice was trained using some of the same data used to build S-CAP, a random train and test split of the pathogenic data would have resulted in the inclusion of variants in the test set that were used to train the MutPred Splice classifier. To ensure zero information leakage between the training and testing datasets, we carefully built a test set that excluded all MutPred Splice training data.

### Recessive v. Dominant Classifiers

We developed separate classifiers for recessive and dominant acting variants in the 3’ and 5’ core splice site regions. We opted not to include dominant and recessive classifiers for the other 4 regions as we did not have a sufficient number of pathogenic variants to train and evaluate multiple models and, in our exploration, they made less of an impact. Intuitively, for core variants the molecular phenotype is obvious, a loss of splicing. This places the full burden on predicting if this change will result in disease, a task heavily dependent upon whether the variant acts via a recessive or dominant mechanism. Whereas in the other regions, the majority of possible variants have little impact on splicing, making predictions as to whether or not the variant will have an effect on splicing is the primary challenge, a task unrelated to inheritance mode.

When evaluating a patient, core variants observed as heterozygous are routed to the dominant classifier and variants observed to be homozygous are routed to the recessive classifier. Since there is no genotype information available for the labeled pathogenic and benign variants, we developed a framework to label variants as dominant or recessive based on their occurrence in a control population. We labeled pathogenic variants observed as heterozygous in the control population as likely recessive, because a single copy can be harbored with no major issues, and pathogenic variants never observed in the control population as likely dominant. Benign heterozygous variants that are never observed as homozygous in the healthy control population provide little information regarding their potential to cause a recessive acting disease. In this case, we tagged benign heterozygous variants never observed to be homozygous as dominant, and benign variants observed as homozygous as recessive. Additionally, as any variant on the X chromosome resembles a homozygous autosomal variant in males, all X chromosome variants were labelled as recessive.

We encoded whether each variant was considered dominant or recessive as a binary feature and trained a single gradient boosting tree model for each region. Then, we found two 95% true positive rate thresholds separately based on test sets of only dominant-tagged and only recessive-tagged variants.

### Patient Datasets

Sequencing and diagnosis for all patients was performed by other laboratories. All patient data were downloaded by requesting access to the European Genome-Phenome Archive (EGA) and the database of Genotype and Phenotype (dbGaP) databases. Variant call files (VCFs) for 11 patients were submitted by the Deciphering Developmental Disorders (DDD) study in the European Genome-Phenome Archive (EGA) study EGAS00001000775. An additional 3 patient VCFs were submitted to the database of Genotype and Phenotype (dbGaP) study phs000655.v3.p1.

## Data availability

S-CAP scores for all rare variants in the predefined splicing region in the human genome, along with the source code repository and final trained models for the S-CAP classifier, will be made available via the S-CAP website upon publication.

## URLs

S-CAP website, http://bejerano.stanford.edu/scap (upon publication)

S-CAP codebase, https://bitbucket.org/bejerano/splicing_classifier (upon publication)

RVIS, http://genic-intolerance.org/data/RVIS_Unpublished_ExACv2_March2017.txt

SPIDEX, http://www.openbioinformatics.org/annovar/spidex_download_form.php

LINSIGHT, http://compgen.cshl.edu/˜yihuang/LINSIGHT/

Haploinsufficiency, https://doi.org/10.1371/journal.pgen.1001154

LabranchOR, http://bejerano.stanford.edu/labranchor/

ExAC, ftp.broadinstitute.org/pub/ExAC_release/release1/regional_missense_constraint/

## Acknowledgements

We thank Ron Dror, Aaron Wenger, Mark Berger, Johannes Birgmeier and all members of the Bejerano Lab for useful discussions, feedback and advice. P.D.S and D.N.C receive financial support from Qiagen through a license agreement with Cardiff University. This work was funded in part by Stanford Graduate Fellowship and the Computational and Evolutionary Genomics Fellowship to K.A.J. and the Stanford Pediatrics Department, DARPA, a Packard Foundation Fellowship and a Microsoft Faculty Fellowship to G.B.

## Author Contributions

K.A.J., J.M.P. and G.B. designed the study. K.A.J., J.M.P. and J.S.Y. developed the features, trained the model and evaluated the results. P.D.S. and D.N.C. curated the HGMD data. J.A.B. provided patient exome cases and feedback. K.A.J, J.M.P. and G.B. wrote the manuscript. All authors reviewed the manuscript.

## Supplementary Figures

**Supplementary Table 1.** Description of features used to build S-CAP. The chromosome, gene, exon and variant level features were used in all models for all regions. Additional features were specific to certain regions. These are enumerated in the rows below the variant level features section.

| Feature Category | Features | Description |
| --- | --- | --- |
| Chromosome Level | X, Y or not XY | 3 binary features to indicate whether a variant is found on the autosomes or the X or Y chromosome |
| Gene Level | RVIS | Measure of gene mutability. Observed v. Expected variant abundance |
|  | pLI | Measure of gene loss of function tolerance. |
|  | haploinsufficiency score | Recessive or dominant gene inheritance |
| Exon Level | # rare 3' core variants | rare variants observed in 3' core region |
|  | # rare 5' core variants | rare variants observed in 5' core region |
|  | # common 3' core variants | common variants observed in 3' core region |
|  | # common 5' core variants | common variants observed in 5' core region |
|  | # rare 3' extended variants | rare variants observed in 3' extended region |
|  | # rare 5' extended variants | rare variants observed in 5' extended region |
|  | # common 3' extended variants | common variants observed in 3' extended region |
|  | # common 5' extended variants | common variants observed in 5' extended region |
|  | exon % identity | Measure the % identity of hg19 exon with each of the 99 species in the multiz100way. Take top 6 PCA components after fitting a PCA to all coding exons. |
|  | exon length | Number of bases in the exon |
|  | exon length % 3 | Number of bases modulo 3 |
|  | MPC | Regional mutational constraint score |
| Variant Level | Spectrum Kernel | Count of all 3-mers introduced and removed by mutation |
|  | MaxEntScan | The difference in motif match |
| | SPIDEX | Predicted $\Delta\psi$ (change in exon expression) |
|  | distance to 5' SS | Number of bases to the 5' splice site |
|  | distance to 3' SS | Number of bases to the 3' splice site |
|  | CADD | Functional data SVM-based classifier |
|  | LINSIGHT | Conservation based model to identify variants under negative selection. |
|  | PhyloP | Base-pair conservation across primates, mammals and vertebrates |
|  | PhastCons | Regional conservation across primates, mammals and vertebrates |
| 3' Intronic | LaBranchoR | Sequence based deep learning branchpoint prediction |
| 3' Intronic, 3' Core, Exonic | 3' cryptic splice site creation terms | MaxEntScan based feature to measure cryptic splice creation near the 3' side |
| Exonic, 5' Core, 5' Extended, 5' Intronic | 5' cryptic splice site creation terms | MaxEntScan based feature to measure cryptic splice creation near the 5' side |
| 3' Core, 5' Core | Zygosity | Indicate whether the variant is seen in a heterozygous or homozygous state. |

**Supplementary Figure 1.**
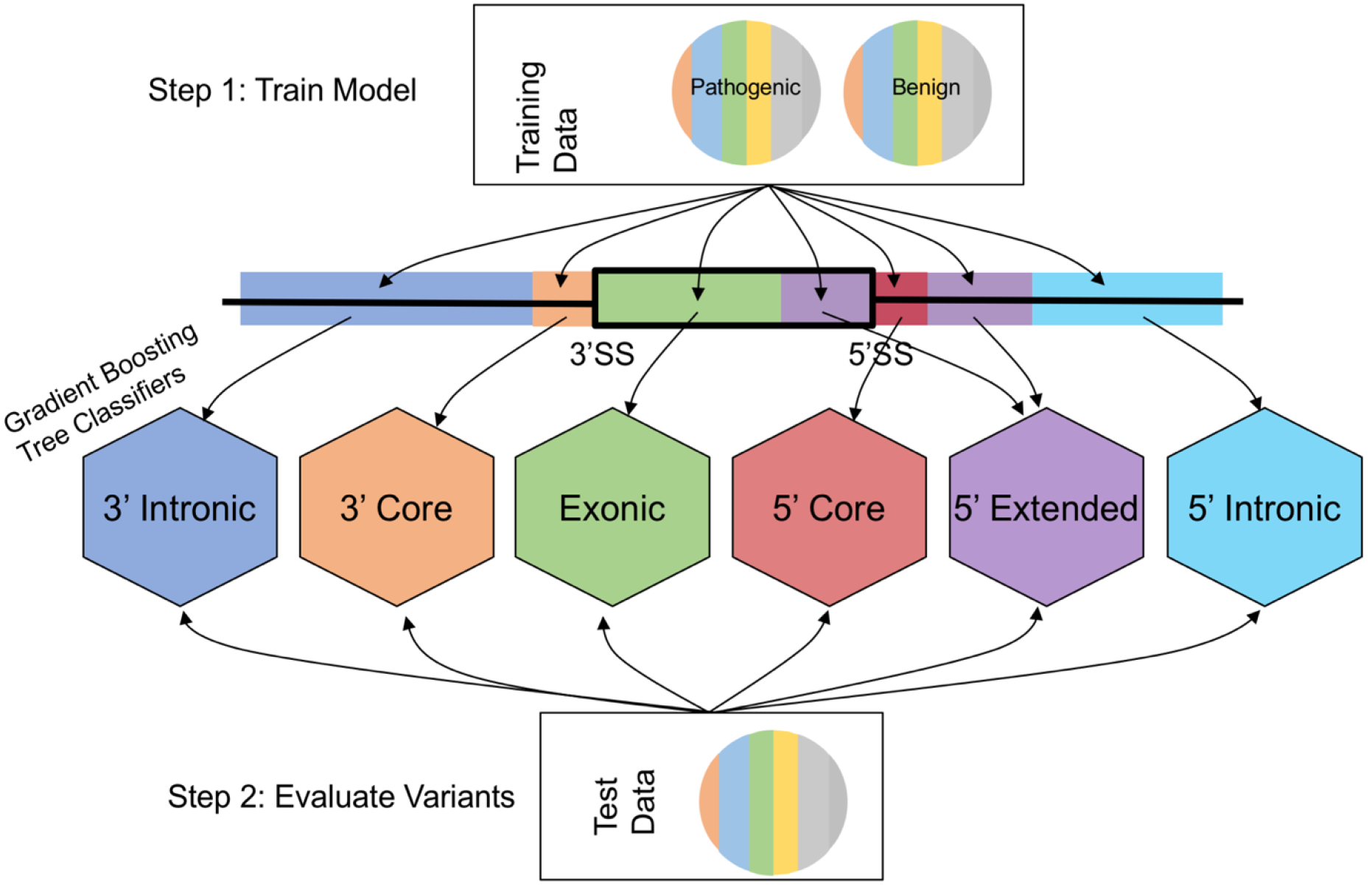
Framework for training and evaluating 6 pathogenicity models. The splicing region is split into 6 independent regions as defined in Fig. 1c, and a separate model is trained for variants residing in each region. Given a set of variants to be scored, we calculate the S-CAP score for each variant by using the corresponding model associated with the region where the variant is found.

**Supplementary Figure 2.**
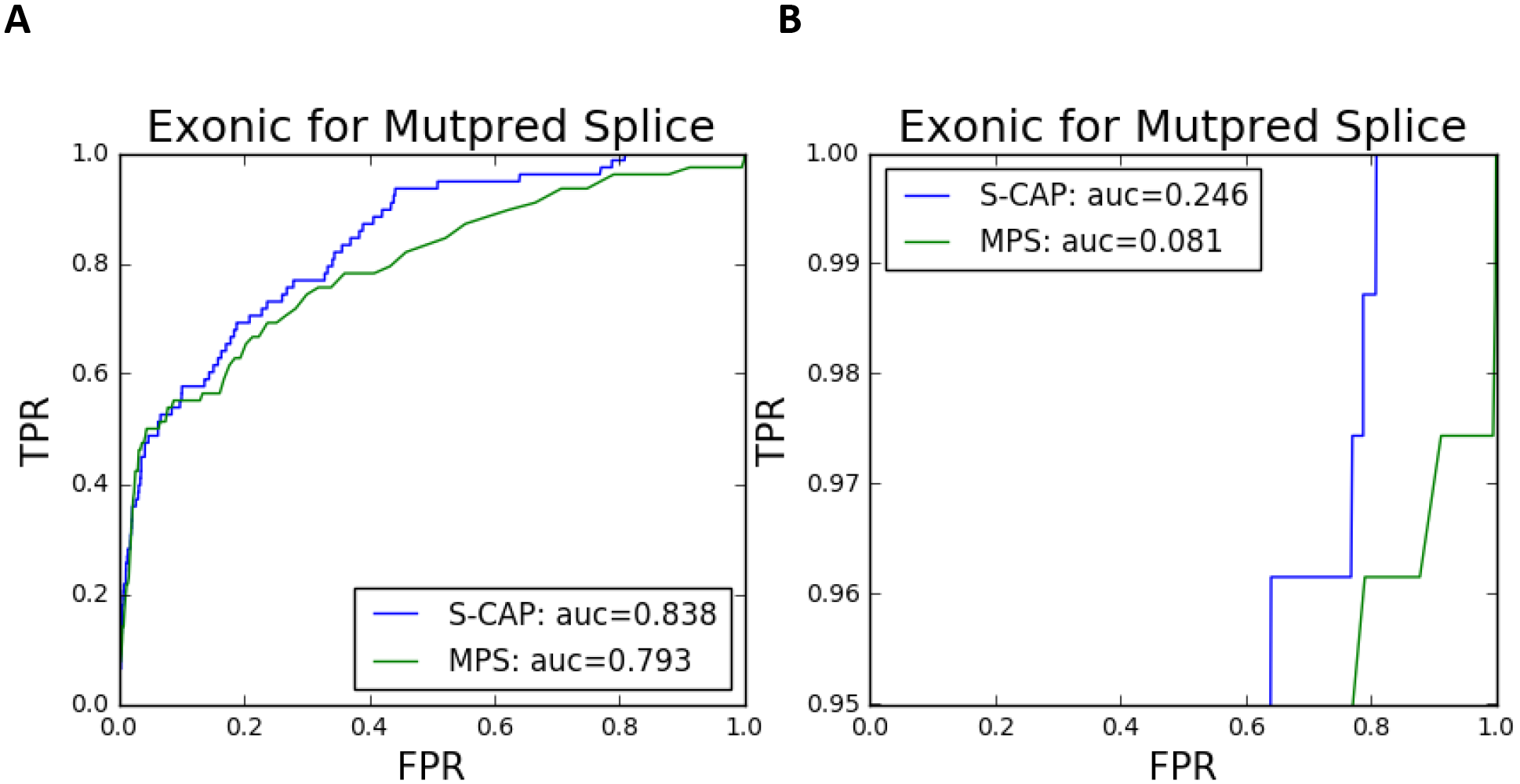
Performance of S-CAP compared to MutPred Splice. MutPred Splice is a computational method for predicting the pathogenicity of exonic synonymous variants. MutPred Splice was trained by its authors using a subset of the pathogenic data used to train/test S-CAP. As a result, we need to independently test MutPred Splice on a set of variants that was not used in its training. This test set comprises rare synonymous variants from HGMD added to the database in 2013 or later. On this set S-CAP improves on MutPred Splice by 5.7% when comparing the overall AUC. S-CAP performs especially well in the high sensitivity domain and improves on the MutPred Splice hsr-AUC by 204%.

**Supplementary Figure 3.**
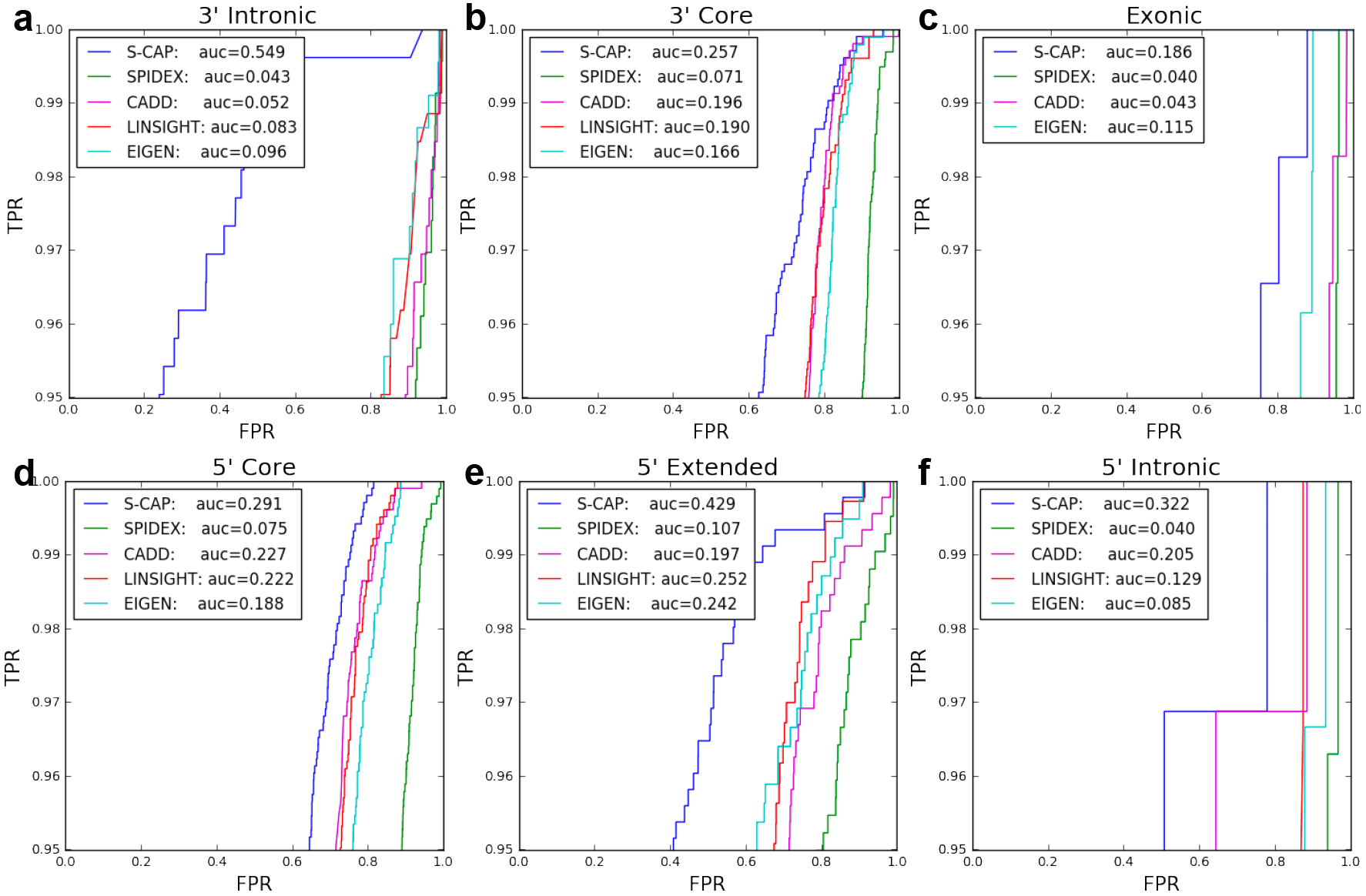
S-CAP performance in the high sensitivity region. The hsr-AUC curve is formed by subsetting the overall AUC to just the region where pathogenic variants are correctly classified over 95% of the time. An hsr-AUC curve is calculated for each of the regions as defined in **Fig. 1c**. S-CAP improves on the next best method’s hsr-AUC by 472% in the 3’ intronic region (A), 31.1% in the 3’ core sites (B), 61.7% in the exonic region (C), 0.28.2% in the 5’ core sites (D), 70.2% in the 5’ extended region (E) and 0.57.1% in the 5’ intronic region (F). None of the existing methods consistently outperforms the others.

**Supplementary Figure 4.**
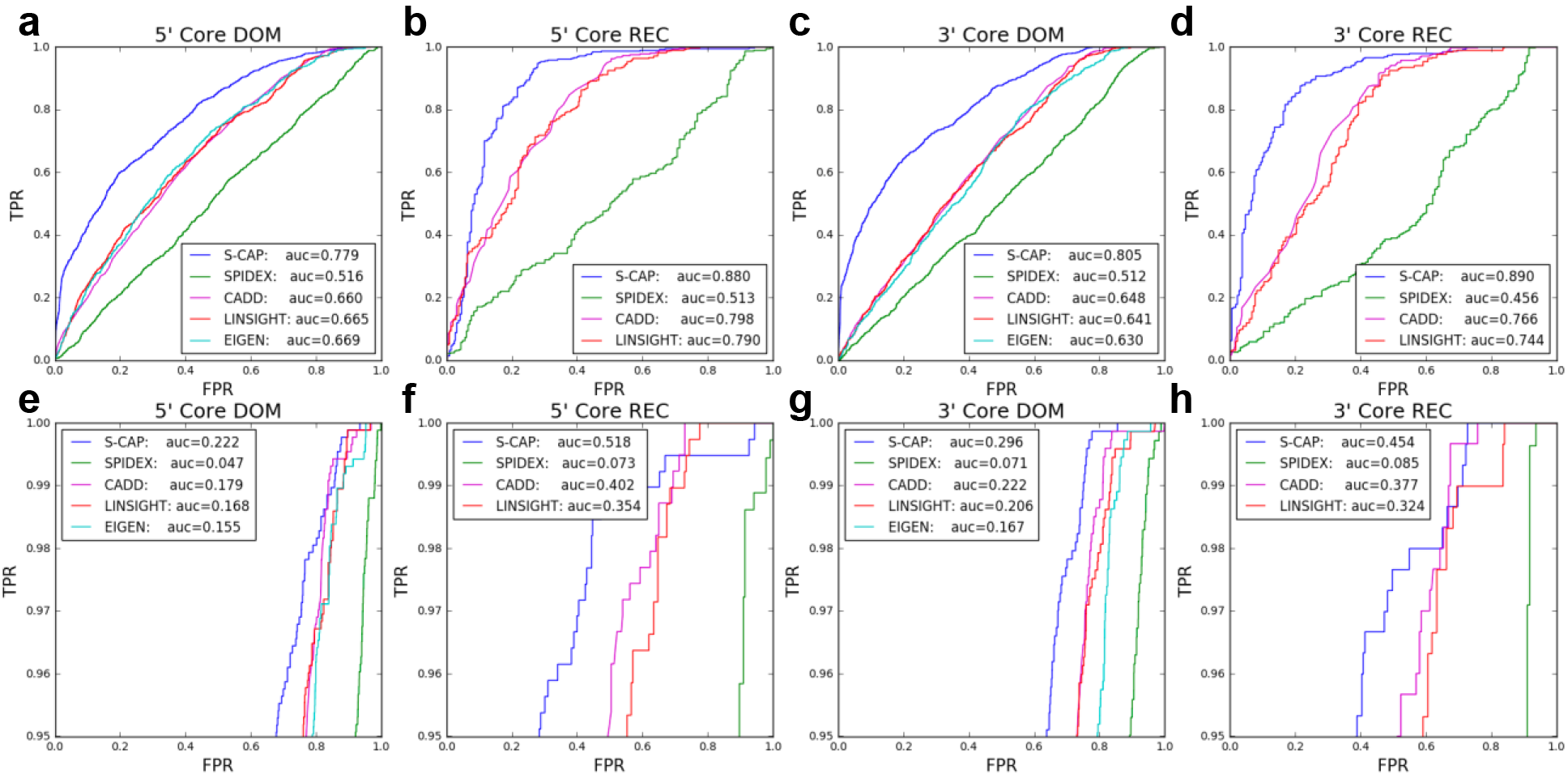
Performance on recessive and dominant classes. The distribution of the underlying features is dramatically different for dominant and recessive variants. This results in a big difference in performance when classifying recessive and benign variants in the core splice site regions. S-CAP achieves an AUC (A) of 0.779 on dominant tagged variants and (B) of 0.880 on recessive tagged variants in the 5’ core region. There is a similar performance difference in the 3’ core region where S-CAP achieves an AUC (C) of 0.805 on dominant tagged variants (D) and of 0.890 on recessive tagged variants. In the high sensitivity region, S-CAP achieves an hsr-AUC (E, F) of 0.222 on dominant tagged variants and of 0.518 on recessive tagged variants in the 5’ core region (G, H) and of 0.296 on dominant tagged variants and of 0.454 of recessive tagged variants in the 3’ core region. S-CAP AUC and hsr-AUC are consistently better than those of all other tools, and no existing method consistently outperforms the others.

